# Functional analysis of the polar amino acid in the TatA transmembrane helix

**DOI:** 10.1101/2021.11.30.470661

**Authors:** Binhan Hao, Wenjie Zhou, Steven M. Theg

## Abstract

The twin-arginine translocation (Tat) pathway utilizes the proton-motive force (pmf) to transport folded proteins across cytoplasmic membranes in bacteria and archaea, as well as across the thylakoid membrane in plants and the inner membrane in mitochondria. In most species, the minimal components required for Tat activity consist of three subunits, TatA, TatB, and TatC. Previous studies have shown that a polar amino acid is present at the N-terminus of the TatA transmembrane helix (TMH) across many different species. In order to systematically assess the functional importance of this polar amino acid in the TatA TMH in *Escherichia coli*, a complete set of 19-amino-acid substitutions was examined. Unexpectedly, although being preferred overall, our experiments suggest that the polar amino acid is not necessary for a functional TatA. Hydrophobicity and helix stabilizing properties of this polar amino acid were found to be highly correlated with the Tat activity. Specifically, change in charge status of the amino acid side chain due to pH resulted in a shift in hydrophobicity, which was demonstrated to impact the Tat transport activity. Furthermore, a four-residue motif at the N-terminus of the TatA TMH was identified by sequence alignment. Using a biochemical approach, the N-terminal motif was found to be functionally significant, with evidence indicating a potential role in the preference for utilizing different pmf components. Taken together, these findings yield new insights into the functionality of TatA and its potential role in the Tat transport mechanism.

## Introduction

The twin-arginine translocation (Tat) pathway, a protein transport machinery, is found in bacteria, archaebacteria, chloroplasts, and plant mitochondria. In bacteria, the Tat pathway is involved in many critical biological processes, including cell division, stress tolerance, and electron transport (Berks, 2015; Gohlke et al., 2005; Ize et al., 2003). In plants, the Tat pathway transports several proteins that are essential for photosynthesis across thylakoid membranes (Clark and Theg, 1997). In haloarchaea which normally possess high cytoplasmic salt concentration leading to faster protein folding, the Tat pathway serves almost 50% of secretome (Ghosh et al., 2019). Unlike the similarly ubiquitous Sec pathway, the Tat pathway is able to transport folded proteins (Clark and Theg, 1997) while requiring only the proton motive force (pmf), with no contribution from NTP hydrolysis (Braun et al., 2007).

The Tat system in most species consists of three subunits, TatA, TatB, and TatC (Berks, 2015). It has been shown that all three subunits can form a complex which works as a receptor for signal peptides of Tat substrates. After the receptor complex binds to the Tat substrate, more TatA subunits are recruited to the translocon. While TatB and TatC are present in a 1:1 stoichiometry in the complex, TatA is present at much higher stoichiometries (8-40 folds) (Leake et al., 2008; Mori et al., 2001; Sargent et al., 2001).

The structures of the three individual Tat subunits have been determined (Hu et al., 2010; Pettersson et al., 2018; Rollauer et al., 2012; Zhang et al., 2014). Both TatA and TatB possess a short transmembrane helix (TMH) followed by a hinge region, one or more amphipathic helices (APH), and then a relatively unstructured C-terminus. Conversely, TatC has six TMHs and exhibits a cupped hand shape. Remarkably, the Tat machinery transports folded proteins of different sizes without introducing significant ion leakage (Asher and Theg, 2021; Teter and Theg, 1998). Recent work suggests that this occurs in the absence of channel/pore like structure, unlike the well-characterized Sec translocon (Tsirigotaki et al., 2017). It has been proposed that Tat pathway operates through lipid-lined toroidal pores formed in a destabilized membrane, rather than common proteinaceous pore (Asher and Theg, 2021; Brüser and Sanders, 2003; Hao et al., 2021).

Previous studies have demonstrated that TatAs have a highly conserved short TMH (Hao et al., 2021) which could cause membrane destabilization and is critical to the overall Tat function (Hao et al., 2021; Hou et al., 2018). Such short TMHs make the TatA protein energetically unfavorable to span on the membrane bilayer. Furthermore, it has been shown that almost all TatA family proteins contain a polar residue at the N-terminus of its TMH (Hicks et al., 2003). In *Escherichia coli* (*E. coli*), TatA has a glutamine (Q) at the 8^th^ position (TMH: 6^th^ – 20^th^), while in pea chloroplasts, TatA has a glutamic acid (E) instead at the corresponding position. A cysteine-scanning study in *E. coli* TatA illustrated that cysteine substitution for the 8^th^ glutamine (Q8C mutant) led to the complete loss of Tat activity, whereas cysteine substitutions for all the other residues in the TMH retained Tat transport activity similar to that of the wtTatA (Greene et al., 2007). Further studies showed that *E. coli* TatA asparagine substitution (Q8N) and Q8E mutants retained the ability to export TMAO (trimethylamine *N*-oxide) reductase (TorA) to the periplasm (Greene et al., 2007), while the alanine (Q8A) mutant was not able to transport either TorA (Greene et al., 2007) or CueO (Alcock et al., 2016). However, another study showed a somewhat conflicting result that *E. coli* Q8A and Q8C mutants retained TorA transport capability when overexpressed (Warren et al., 2009). On the other hand, in pea or maize chloroplasts, it was shown that all of the 10^th^ glutamic acid substitution variants, E10A, E10D, and E10Q, severely affected Tat transport in a biochemical complementation assay (Dabney-Smith et al., 2003; Fincher et al., 2003). Interestingly, although the chloroplast TatA E10Q mutant exhibited extremely low Tat activity, the *E. coli* TatA Q8E mutant displayed high Tat transport activity, suggesting that residue preference was not identical for different species. Such different preference was also shown with chimeric pea/*E. coli* TatA derivatives (Hauer et al., 2017). Furthermore, the *Bacillus subtilis* Tat system contains two sets of TatA/TatC subunits (TatAd/TatCd and TatAy/TatCy), and those TatAs actually lack this “conserved” polar residue in the TMH. Although extensive effort has been made to investigate the function of this “conserved” polar amino acid in TatA, the lack of a complete set of mutants and use of different transport assays has allowed the role of this residue at the analogous position to remain an open question.

In this study, we analyzed the amino acid requirement for the 8^th^ residue in *E. coli* TatA through a complete set of Q8 mutants. Both qualitative and quantitative assays were conducted to compare the activity between TatA mutants. Unexpectedly, we found that polar amino acids were preferred overall but actually not necessary for a functional TatA. However, the Tat activity was highly correlated with hydrophobicity and helix stability of the 8^th^ residue in TatA. Furthermore, by sequence alignment and biochemical analysis, a four-residue motif at the N-terminus of the TatA TMH was found to be functionally significant. These results expand our understanding of the potential function of TatA and provide new insight into the Tat transport mechanism.

## Results

### 1. Polar amino acids at the 8^th^ position in *E. coli* TatA TMH are preferred but not necessary for minimal Tat activity

To systematically assess the characteristic requirements of the 8^th^ residue in *E. coli* TatA, Q8 was replaced by all the other 19 amino acids and their effects on Tat activity was measured. Membrane stability of TatA Q8 mutants was assessed by membrane isolation followed by carbonate wash, and all mutants displayed an integrated presence in the membrane (Supplemental figure 1). Minimal Tat activity was subsequently screened by SDS growth assays (Ize et al., 2003) for each of the TatA Q8 mutants. All polar amino acid substitutions (Q8N, Q8S, Q8T, Q8D, Q8E, Q8H, Q8K, and Q8R) exhibited sufficient Tat activity to grow in LB medium containing 10% SDS (Figure 1A). Surprisingly, TatA Q8A, Q8C, Q8G, and Q8P mutants, whose substituted amino acids do not possess strong polar side chains, also showed a growth activity in the same medium (Figure 1B, C). The rest of substitutions (Q8F, Q8I, Q8L, Q8M, Q8V, Q8W, and Q8Y) displayed no growth activity in LB medium containing 5% SDS, indicating a complete loss of Tat activity (Figure 1C). An *in vivo* transport assay was implemented to further confirm the SDS growth assay result. In this assay, a FLAG-tagged *E. coli* native Tat substrate, SufI-FLAG, was overexpressed in the TatA Q8 mutants. After 2.5 hours transport at 37°C, a periplasmic fraction was isolated by treated with EDTA/lysozyme and osmatic shock (Petiti et al., 2017a), a harsh method that minimizes the loss of periplasmic proteins. The precursor and mature forms of SufI-FLAG were separated by SDS-PAGE according to their size. This assay revealed that, despite their ability to grow in 10% SDS, the Q8A, Q8C, Q8D, Q8E, Q8G, Q8P, Q8R, and Q8T mutants displayed significantly lower Tat activity (<25%) compared to wild-type TatA (Figure 1D).

**Figure 1.**
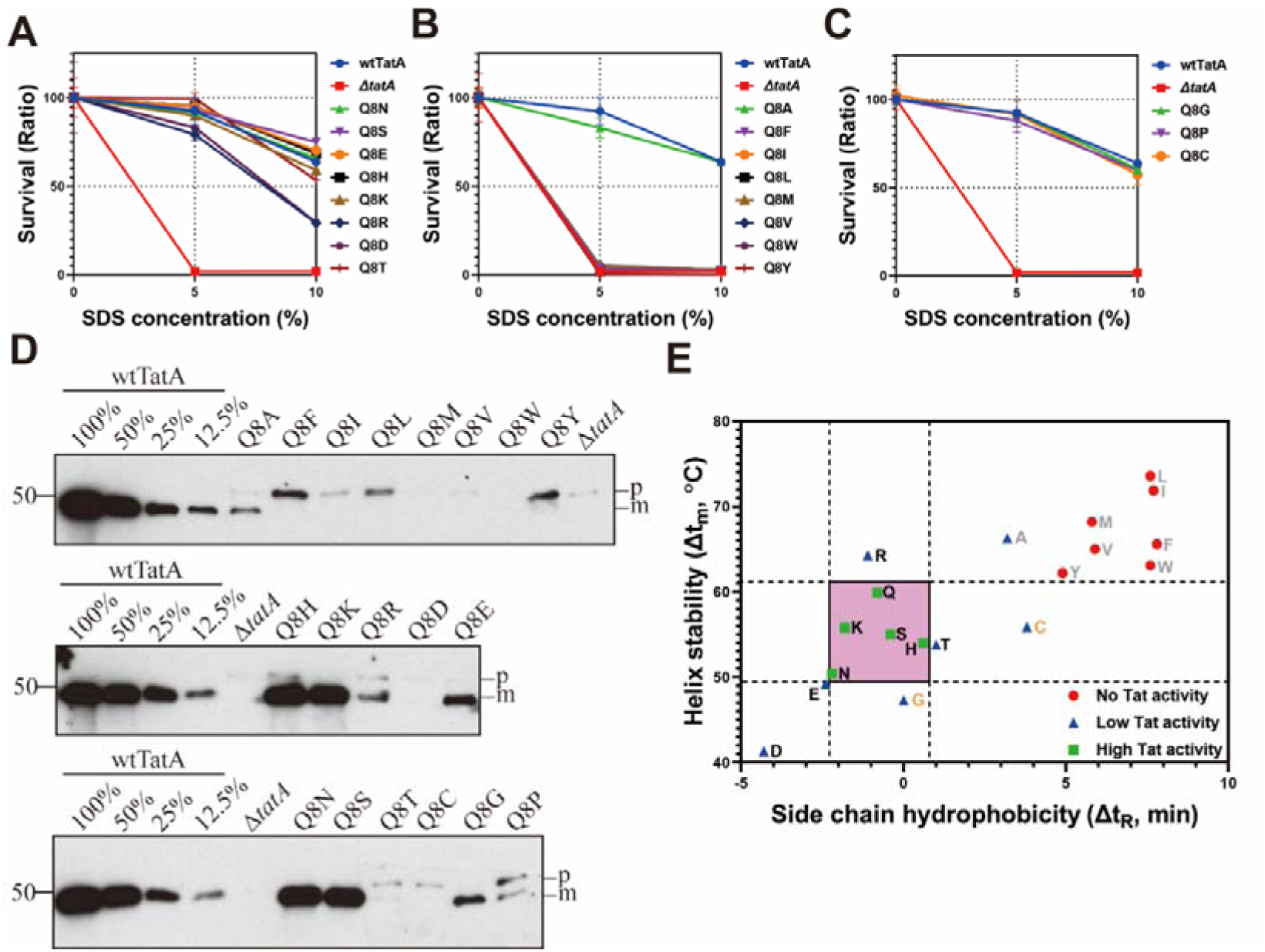
Polar residues at the eighth position of the *E. coli* TatA TMH are preferred but not necessary for minimal Tat activity. (A-C) Minimal Tat transport abilities of the TatA Q8 variants were evaluated by the growth performance in LB medium with indicated SDS concentrations. wtTatA served as the positive control, while Δ*tatA* acted as the negative control. Survival ratio denotes the ratio of the OD_600_ of the cells grown in the presence of the corresponding SDS concentration to the OD_600_ of the cells grown without SDS. Three biological replicates for each mutant were included in the experiments. (D) Western blot results of the *in vivo* transport assays with the TatA Q8 variants. Dilutions of the amount of transported SufI-FLAG in the wtTat cells were shown for the comparison of the transport performance in the TatA Q8 variants. p, precursor and m, mature. (E) Plot of helix stability (Δt_m_) versus side chain hydrophobicity (Δt_R_) of amino acids colored according to their activity in the 8^th^ position of TatA. Different levels of the Tat transport activities are denoted with the following rules: no Tat activity, no growth in the LB medium containing 5% SDS; low Tat activity, mutants who could grow in the LB with SDS but showed less than 50% transport activity than wtTat in the *in vivo* transport assay in (D); high Tat activity, mutants with more than 50% of the wtTat activity in (D). Identities of the amino acids with high Tat activity were clustered in the ranges of moderate helix stability and side chain hydrophobicity, which is boxed in pink.

Pulse-chase assays, representing a more fine-grained assessment of Tat transport activity (Palmer et al., 2010), were conducted with all the functional TatA Q8 mutants to compare their transport rates. Figure 2A shows the quantitative analysis of the pulse chase results (Supplemental figure 2). An exponential plateau model *y* = *y*_max_ – (*y*_max_ – *y*_0_) × *e*^−*kt*^ was applied in data fitting. Estimated initial transport rate (*V*_0_) derived from the fitting curve was used to compare the transport activity between the TatA Q8 mutants (Figure 2B). Unexpectedly, the Q8H mutant exhibited the highest *V*_0_ (228% of the *V*_0_[wtTatA]). Q8N, Q8S, and Q8E variants of TatA, exhibited 58%, 37% and 33% of the *V*_0_[wtTatA], respectively. However, Q8A, Q8C, Q8D, Q8G, Q8K, Q8P and Q8R showed relatively lower transport rate (less than 25% of *V*_0_[wtTatA]). One unanticipated finding was that the Q8T, Q8D and Q8R mutants, whose 8^th^ glutamine was replaced by a polar amino acid, showed even lower transport rates than the poorly performing Q8A mutant. The results from pulse-chase assays were overall consistent with the *in vivo* transport assay except that Q8K performed relatively worse in the pulse-chase assay than in the *in vivo* transport assay.

**Figure 2.**
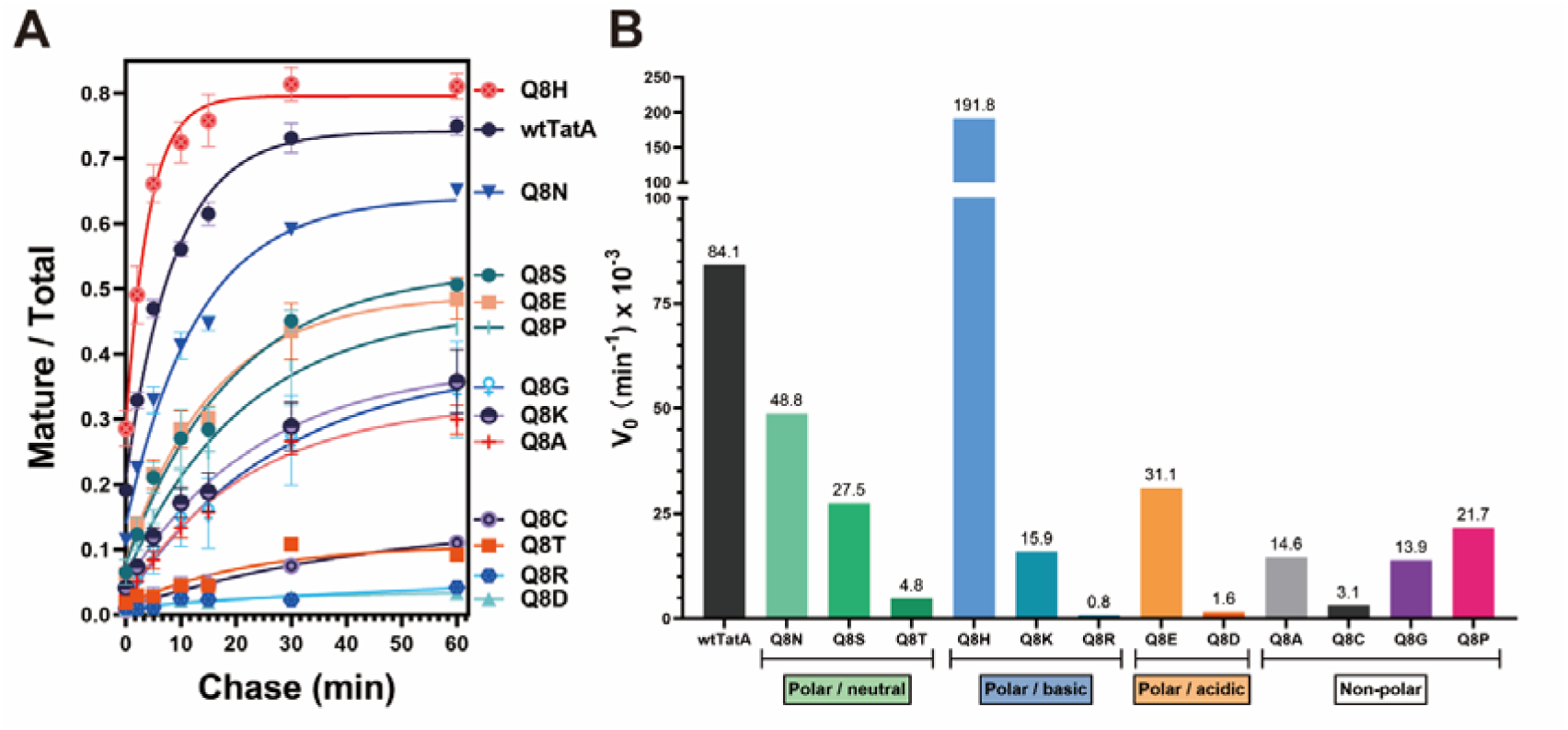
Quantitative analyses of the Tat activity in the TatA Q8 variants. (A) Plot of the transport kinetics of the TatA Q8 variants in pulse-chase experiments. Identities of wtTatA and TatA Q8 variants are shown in the right. Mature/total denotes the ratio of the mature band to the sum of the precursor and mature bands revealed by autoradiography of the samples from the pulse-chase experiment transporting SufI; chase time is denoted on the abscissa. Lines show best fits of an exponential model to the points arising from at least two biological replicates for each mutant. (B) Plot of the initial velocities (V_0_) in the TatA Q8 variants transporting SufI. Exact values of the V_0_ are shown above the bars. TatA Q8 variants are grouped based classification of the amino acids.

In summary, polar amino acids are overall strongly favored at the 8^th^ position in the *E. coli* TatA but not necessary for a functional Tat translocon. Moreover, not all polar amino acid substitutions showed higher transport ability than non-polar substitutions.

### 2. The Tat activity of Q8 mutants is correlated with the side chain hydrophobicity and helix stabilizing properties of the substituted amino acid

Our data clearly show that polar amino acids are preferred overall in the TatA 8^th^ position, but are not strictly necessary to retain Tat activity (ex. Q8A, Q8C, Q8G, Q8P). Moreover, some of the polar residue substitutions (i.e., Q8D, Q8R and Q8T), even though they retained the minimal Tat activity required to grow in LB medium with SDS, displayed significantly lowered transport activity. Such results could not simply be explained by the binary categories, polar or non-polar, of amino acids. In order to understand the functional importance of this position in the *E. coli* TatA, a further analysis of the amino acids’ preference is needed. On one hand, previous studies showed that the overall hydrophobicity of the TatA TMH might affect TatA function in thylakoids (Hauer et al., 2017). On the other, a recent study has demonstrated that the hydrophobic mismatch between the TatA TMH and membrane bilayer is critical to Tat function (Hao et al., 2021; Mehner-Breitfeld et al., 2021). To maintain such a hydrophobic mismatch, a relatively stable alpha helix structure is likely to be required. To assess these properties, the hydrophobicity and α-helix stability of each amino acid (Monera et al., 1995) were plotted together with the transport performance of the corresponding TatA Q8 mutants (Figure 1E). Interestingly, a high correlation between Tat activity and the combination of hydrophobicity and α-helix stability was observed: 1) the Q8 mutants which completely lost Tat activity clustered at the top-right corner, suggesting that very high hydrophobicity and helix stability at this position are detrimental for Tat transport. 2) all mutants with high Tat activity were clustered in the middle ranges of hydrophobicity and helix stability (Figure 1E, pink box). These observations draw our attention to the importance of a moderate hydrophobicity and α-helix stability of the 8^th^ amino acid in *E. coli* TatA.

### 3. Charge status of the side chain of the polar amino acid affects the TatA function

Due to the nature of polar amino acids with electrically charged side chains, previous studies have reported that the hydrophobicity of those amino acids changes dramatically under different pH conditions (Kovacs et al., 2006; Mant and Hodges, 2002). For instance, the side chain of histidine (pKa_(His)_ = 6.04) is relatively hydrophobic (similar as Ala) at pH 7 (mostly uncharged), while it becomes one of the most hydrophilic side chains when the pH is lower than 5 (mostly charged). In this study, we hypothesized that a moderately hydrophobic side chain is required for the 8^th^ residue in the *E. coli* TatA. To further investigate this hypothesis, a comparison of Tat transport rates between wtTatA and Q8H was studied under different pH conditions. The pulse-chase experiments in Figure 2 showed that the Q8H mutant performed approximately 2-fold better than the wild-type TatA at pH = 7.0 where most of (90%) the histidines in TatAs were presumably deprotonated. To assess whether a change in histidine hydrophobicity would alter activity of the Q8H mutant, we measured transport rates by pulse-chase at pH 5.0, 6.0 and 8.0 (Figure 3A). Figure 3 shows that wtTatA and Q8H had different behaviors when transporting SufI at different pH. First, wtTatA and Q8H mutant exhibited optimal transport rates at different pH values (Figure 3B and C). wtTatA displayed optimal transport activity at pH 5.0, while the optimal pH for Q8H mutant was 7.0. However, their highest V_0_ values were comparable to each other (Figure 3D). Second, *V*_0_[wtTatA] was significantly higher than *V*_0_[Q8H] at pH = 5.0 (Figure 3E) whilst *V*_0_[wtTatA] was significantly lower than *V*_0_[Q8H] at pH = 7.0 (Figure 3G). Third, wtTatA had a very similar *V*_0_ as Q8H mutant at pH = 8.0 (Figure 3H) where 99% of the histidines were presumably deprotonated, a similar status as uncharged glutamine in wtTatA. In summary, the charged status of the histidine side chain, a model residue for comparison with the wtTatA, affects its transport.

**Figure 3.**
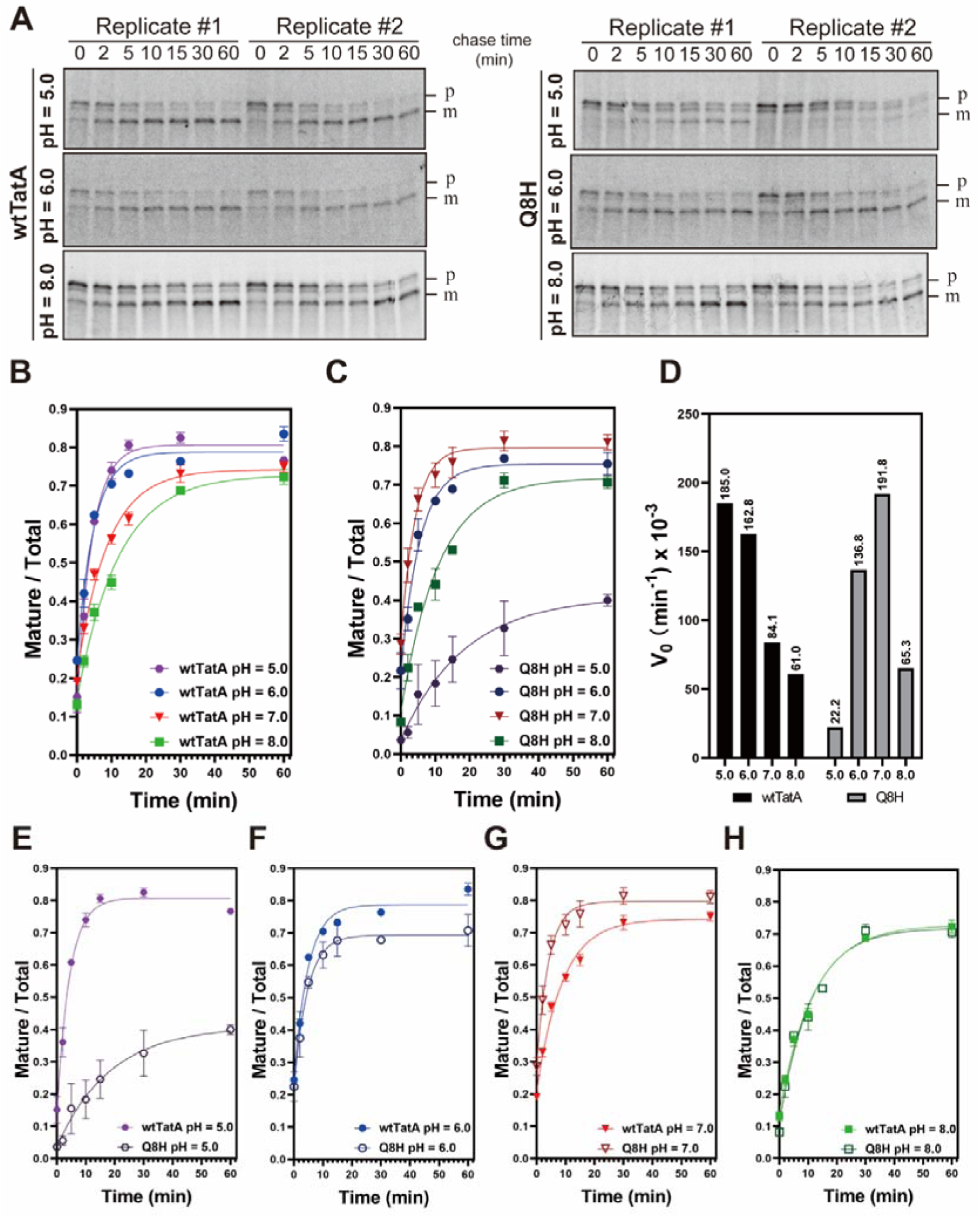
Comparison of the Tat activity of wtTat and Q8H mutant at different pH conditions. (A) Radioautography of the wtTatA and Q8H variants transporting SufI at different pHs. Samples from the pulse-chase experiment were subjected to SDS-PAGE. 8-16% polyacrylamide gels were used to separate the mature SufI (m) from the precursor (p). Corresponding times of the chase steps are shown on top. Two replicates for each variant were included in this experiment. (B-C) Quantitative analyses of the transport performance of the wtTatA (B) and Q8H (C) variants based on the radioautography in (A). Same analysis approach was used as mentioned earlier. The pH condition in which the experiment was performed is indicated in the legend of the plot. (D) Plot of the initial velocities under different pH conditions between the wtTat and Q8H variants. Initial velocities of the wtTatA under different pH are denoted in black, and initial velocities of the Q8H variant under different pH are denoted in grey. (E-H) Comparison of the transport kinetics between the wtTatA and Q8H variants under the same pH (E, pH = 5.0; F, pH = 6.0; G, pH = 7.0; H, pH = 8.0), with the filled symbols representing the wtTatA, and empty circles representing the Q8H. Other details as described in the legend of Fig. 2.

In the aggregate, our experiments in the pH range 5.0-8.0 are consistent with the hypothesis that a functional Tat translocon requires that its TatA possesses a moderately hydrophobic amino acid at 8^th^ position. In addition, it demonstrates that not only hydrophobicity of different amino acid but also hydrophobicity of same amino acids under different charged status affects its ability to support Tat transport. Together these results provide important insights into understanding the function of the 8^th^ residue in *E. coli* TatA (see discussion).

To further investigate how charged residues in TatA affect the overall Tat transport machinery, we determined how well the membrane can hold a pmf by measuring the ΔpH induced by ATP hydrolysis in inverted membrane vesicles (IMVs) membrane expressing corresponding mutated TatA proteins. According to their charge status at pH 7.0, Q8E, wtTatA, Q8H and Q8K mutants were selected as examples of negatively, neutral, partially positively and completely positively charged residues at the 8^th^ position in *E. coli* TatA (Figure 4A). wtTatA and the Q8E mutant showed similar low membrane leakage outcomes in the presence or absence of the TatBC complex. However, Q8H and Q8K mutants exhibited significant different membrane leakage characteristics when the TatBC complex was present. Q8H, which showed the highest membrane leakage without TatBC, displayed less membrane leakage when TatBC was also expressed on the membrane. In contrast, the Q8K mutant showed the opposite behavior, exhibiting the highest ΔpH without TatBC but developing a dramatically high membrane leak when TatBC was present. That is to say, even though lysine and glutamic acid have similar hydrophobicity at pH 7.0 (Figure 1E), their different charge status might affect how they interact with the TatBC complex, which results in different membrane leakage outcomes. The cause for this behavior is not clear and awaits further investigation.

**Figure 4.**
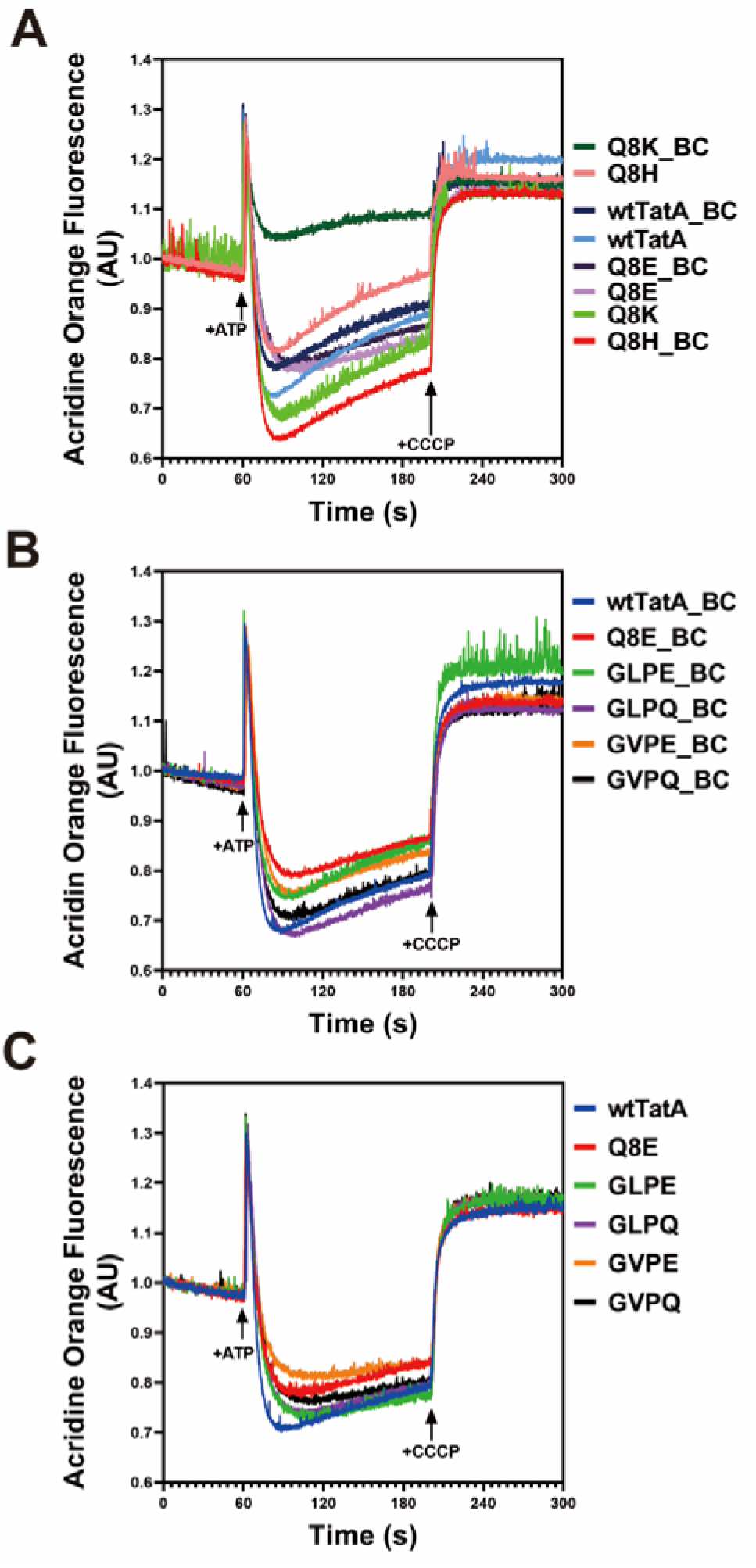
Membrane leakage profiles of TatA mutants. The ΔpH developed across the membrane was measured by the acridine orange quenching in the IMVs. 4 mM ATP was added at the 60 secs to energize the IMV, and 10 μM CCCP was added at 200 secs to dissipate the proton gradient, resulting the recovery of the fluorescence. (A) Measurement of the ΔpH in the DADE (Tat knockout variant) IMVs with indicated overexpressed TatA Q8 variants alone or together with the constitutively expressed TatBC. (B) Measurement of the ΔpH in the DADE (Tat knockout variant) IMVs with indicated overexpressed TatA N-terminal motif variants alone. (C) Measurement of the ΔpH in the DADE (Tat knockout variant) IMVs with indicated overexpressed TatA N-terminal motif variants together with the constitutively expressed TatBC. AU, arbitrary unit.

### 4. Different species have different preferences for polar amino acids in the TatA TMH

Even though in *E. coli* TatA, the polar amino acid in the TMH is the uncharged glutamine, in plant, the Tat system, this position is usually occupied by a glutamic acid, a charged amino acid (Mori and Cline, 2001). Moreover, *Bacillus subtilis*, a gram-positive bacterium, expresses two different kinds of TatA with glycine (TatAc) or serine (TatAy) at the corresponding position. To investigate if such amino acid choices are correlated to species, TatA sequences were aligned according to five categories: proteobacteria (N=7534), actinobacteria and firmicutes (N=6978), chloroplasts (N=242), cyanobacteria (N=149) and archaea (N=151). A conserved 12-residue hydrophobic core was observed (9^th^ – 20^th^ in Figure 5A-E) before the invariant “FG” motif at the end of TMH (Hao et al., 2021). In addition, there is a clear preference for particular amino acids in the TatA TMH at the 8^th^ position with different species (Figure 5F) (Here, we use the numbering from *E. coli* for simplicity. The exact number might be different in different species.) In proteobacteria (representing *E. coli*), histidine (50%) and glutamine (34%) are the top two choices. Interestingly, such choice preferences are consistent with our Tat activity results in that Q8H and wtTatA exhibited the highest Tat transport rates. In contrast, in the other two groups of bacteria, actinobacteria and firmicutes, even though histidine is still the top choice for TatA (46%), glutamine is present in only 2% of the species in these categories, and relatively more of them choose glutamic acid (32%) or glycine (10%) compared to proteobacteria. However, all of chloroplasts, cyanobacteria and archaea TatAs have a highly conserved glutamic acid (>90%) at the 8^th^ position. Remarkably, even though polar amino acids are present in TatA in most species, 1% of actinobacteria and firmicutes (N = 80) and 0.1% of proteobacteria (N = 13) possesses an alanine at that position. This finding further confirms our experimental observations that polar residues at 8^th^ position are not indispensable for a functional TatA (Supplemental Figure 3A and B). Such different preferences in different species suggest that a possible evolutionary pressure exists at this position for the Tat pathway to accommodate various external conditions.

**Figure 5.**
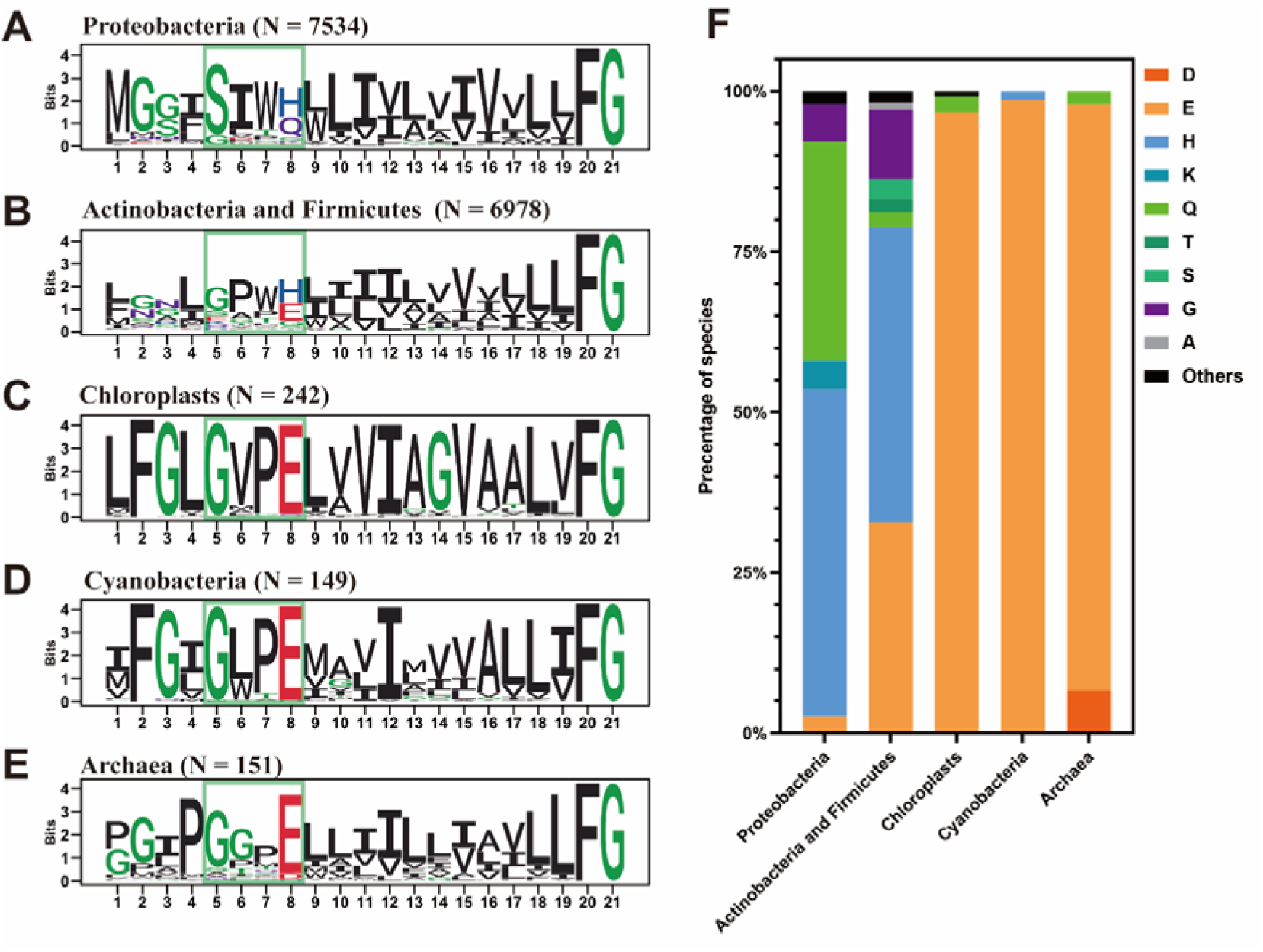
Different species have different preferences at the 8^th^ position in TatA. Sequence logos of the TatA sequences alignment across (A) proteobacteria, (B) actinobacteria and firmicutes, (C) chloroplasts, (D) cyanobacteria, and (E) archaea. Numbers of sequences included in the corresponding categories were denoted as N. The N-terminal motifs were boxed as indicated. (F) Distributions of amino acids selected in the location equivalent to the 8^th^ position of the *E. coli* TatA across different species. Percentage of species with the indicated residues were shown. Amino acids were colored as indicated to the right of the plot.

### 5. A four-residues functional motif at the N-terminus of the TatA TMH is important for Tat activity

Besides the various preference of amino acids at the 8^th^ position in different species, a conserved GxPE motif (the 5^th^ to 8^th^ position in Figure 5C, D and E) was also observed in chloroplasts, cyanobacteria, and archaea TatAs. In contrast, in proteobacteria, actinobacteria and firmicutes, such “GxPE” motif was not obviously presented. Instead, a SIW(H/Q) motif was generally conserved in proteobacteria. To see if these motifs are correlated to different polar amino acids, TatA sequences were further examined based on the identity of the amino acid at the 8^th^ position (Supplemental Figure 3). Supplemental Figure 3A clearly shows that proteobacteria favor “SIWH” and “SIWQ” (i.e., *E. coli*) at the N-terminus of the TatA TMH. In addition, “GLPG”, “SITK” and “S(I/L)WN” motifs were identified in proteobacteria as well. Interestingly, in *E. coli* TatE, “SITK” was at the corresponding location (the 5^th^ to 8^th^ residues). It has been reported that the TatE gene was found only in enterobacteria, a subset of proteobacteria (Yen et al., 2002). Accordingly, in this study, TatA or TatE sequences from enterobacteria were also aligned (Supplemental Figure 3F) to check if such motifs are also present in other TatE genes. Surprisingly, “SIWQ” and “SITK” were extremely conserved across TatA (N=599) and TatE (N=160), respectively, in enterobacteria. Additionally, although “SIWH” was the most conserved motif in proteobacteria, gram-positive bacteria favored “xPWH” in the same position (Supplemental Figure 3B), These results suggest that the four-residues motif at the N-terminus of the TatA TMH might play an important role in TatA function despite minor deviations in different species.

To assess whether the corresponding four-residue motifs are required for specific amino acids at the 8^th^ position to exhibit higher Tat transport activity, we also substituted the 5^th^ to 8^th^ amino acids in *E. coli* TatA according to the above conserved motifs, including “GVPE” (a motif of chloroplasts TatAs), “GLPE” (a motif of cyanobacteria TatAs) and “SITK” (a motif of enterobacteria TatEs). Moreover, to comprehensively understand how the 5^th^ to 7^th^ amino acids affects TatA function, the 5^th^ to 8^th^ residues of *E. coli* TatA were also changed to “GVPQ” or “GLPQ” (Figure 6).

**Figure 6.**
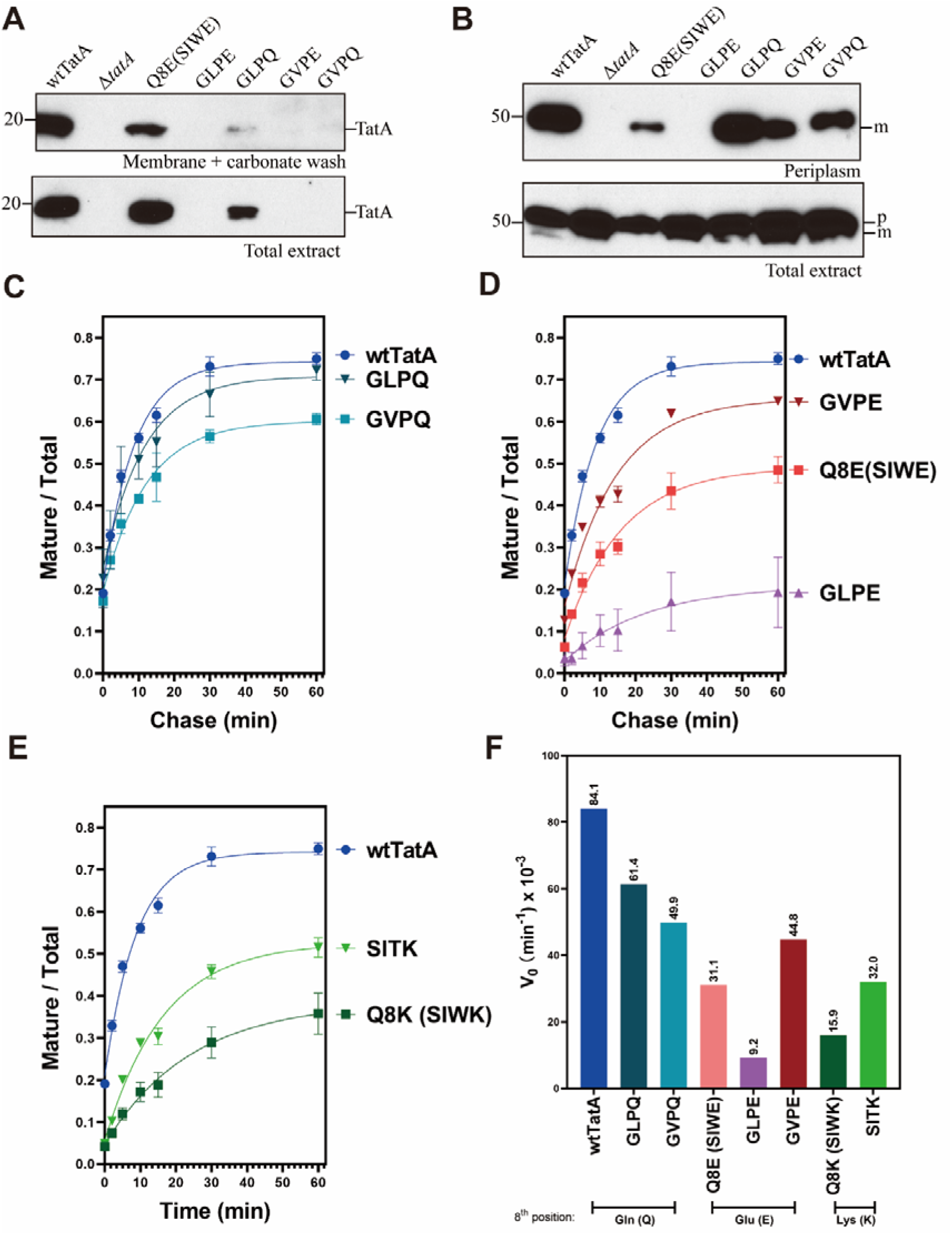
Comparison of the Tat transport activity with the TatA N-terminal motif mutants. (A) Membrane after carbonate washing and total extract samples were subjected to SDS-PAGE and immunoblotting with the α-TatA antibody. Molecular weight is shown on the left. (B) Periplasmic fractions (top) and total extract (bottom) of the cells in the *in vivo* transport assays were isolated. Precursor (p) and mature (m) proteins were labeled on the right, and molecular weight is shown on the left. (C-E) Plots of the transport kinetics for the N-terminal motif mutants in the pulse-chase experiment. At least two replicates were included for each mutant. (F) Plot of the initial velocities (V_0_) of the N-terminal motif mutants in the pulse-chase experiments. V_0_ was obtained from the kinetic plots in (C-E). Same methodology was implemented as described before. Exact values of the initial velocities are displayed on top of bar graph. Other details as described in the legend of Fig. 2.

A recent study has shown that a shorter TMH negatively affects TatA’s membrane stability (Hao et al., 2021). In terms of our new mutants, GVP(E/Q) and GLP(E/Q), it is possible that those mutants lost their ability to stably embed into the membrane since proline (immediately before the 8^th^ residue) usually acts as a helix breaker (Chou and Fasman, 1978) in protein secondary structure. Figure 6A shows that TatA in GVP(E/Q) and GLP(E/Q) mutants exhibited much lower membrane abundance compared to wtTatA or Q8E mutant, indicating that a proline residue severely affects the membrane insertion, even though it did not cause high membrane leakage in those mutants (Figure 4B and C). But, very surprisingly, GVPE mutant displayed a better performance than Q8E in the *in vivo* transport assay (Figure 6B). Pulse-chase assays with those N-motif mutants were also conducted to quantitatively compare their transport rates. Both GLPQ and GVPQ displayed somewhat lower transport rates than wtTatA, suggesting that a disruption of the conserved “SIWQ” motif was deleterious to *E. coli* TatA function (Figure 6C). However, GVPE and SITK mutants exhibited significantly higher transport rates than Q8E and Q8K mutants, respectively, which indicates that the evolutional conserved motifs are functionally more suitable for certain amino acids at the 8^th^ position in TatA (Figure 6D-F). In spite of that replacing to conserved N-motifs were generally beneficial for Tat function, we also observed that the GLPE mutant supported significantly lower transport rates than both the GVPE and Q8E mutants. Such results demonstrate that changes of a single amino acid in the N-terminal motif (even from one hydrophobic residue to another hydrophobic residue) strongly affected the final TatA function. This confirms our hypothesis that the N-terminus of TatA, and not just the well-known polar amino acid, determines the Tat transport activity. Further, it implies that the combinations of these amino acids at the N-terminus of the TatA TMH are potentially tuned to their own membrane environments, and those different motifs could be a strategy to adjust to diverse and subtle biological membrane properties.

### 6. The conserved “GVPE” motif from chloroplast TatA might be responsible for preference of utilization of ΔpH

Even though the GVPE mutant performed significantly better than the Q8E mutant (Figure 6F), it was still less effective than the wild-type SIWQ motif in the *E. coli* TatA.

This result does not point to an explanation of why most chloroplasts and cyanobacteria possess a glutamic acid at the 8^th^ position in their TatAs. It was previously shown that the chloroplast Tat system is highly dependent on ΔpH (Cline et al., 1992), while *E. coli* Tat system does not require ΔpH for active transport (Bageshwar and Musser, 2007a). These energetic preferences are also consistent with their own respective tendencies to partition their pmfs between a ΔpH and a Δψ (Cruz et al., 2001; Zilberstein et al., 1984). To investigate if the invariant motif in chloroplast TatA is related to this energetic preference, IMVs, the bacterial counterpart to chloroplast thylakoids, were made from wtTatA, Q8E and GVPE mutants. A previous study (Bageshwar and Musser, 2007a) used IMVs to test Tat transport ability powered by different components of the pmf. In our hands also, the IMVs made with wild-type *E. coli* Tat displayed a higher transport ability in the presence of nigericin which presumably dissipated the ΔpH. Figure 7A shows that wtTatA and the Q8E mutant displayed consistently higher performance in the presence of nigericin. However, the GVPE mutant exhibited significantly lower transport capability when the ΔpH was dissipated. These results suggest that GVPE mutant relied more on ΔpH for transport than the wtTatA or the Q8E mutant.

**Figure 7.**
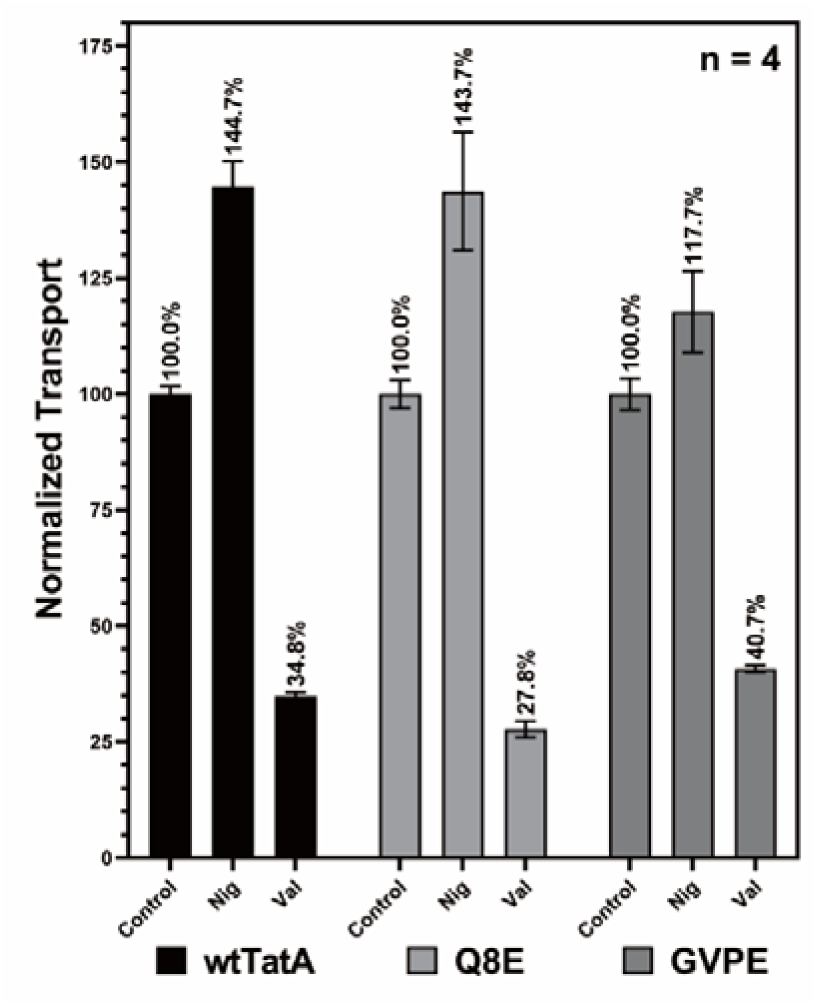
TatA N-terminus mutants display differences in energy utilization preference and thermostability. (A) Comparison of the IMV transport of SufI among wtTatA, TatA Q8E (SIWE), and TatA GVPE mutants in response to 10 μM nigericin or 10 μM valinomycin treatment. The control groups received equivalent amounts of EtOH and were set to 100%. The exact percentage of transport in the experimental groups are shown above the bars. Each treatment group per mutant was performed in four replicates, see Supplemental Figure 4.

## Discussion

An incomplete set of *E. coli* TatA Q8 mutants was a limitation to a full understanding of the function of the “conserved” polar amino acid in the TatA TMH. In this study, we systematically analyzed the Tat transport ability of all the Q8 mutants which the 8^th^ glutamine of *E. coli* TatA was substituted by the other 19 amino acids. Three assays were conducted to compare the transport rates between different mutants, qualitatively and quantitatively. The transport ability results of our TatA Q8 mutants showed that a polar amino acid at the 8^th^ position of TatA is preferred but not necessary for a functional TatA (Figure 1 and 2). Rather, a strong correlation between Tat transport activity and the hydrophobicity and helix stability of the 8^th^ residue was noticed (Figure 1E). It reveals that a residue possessing a moderate range of hydrophobicity and helix stability was preferred for TatA to retain high transport capability. Moreover, by altering the environmental pH in our transport assay, we demonstrated that the Q8H mutant, which contains a protonatable amino acid substitution, exhibited significantly lower transport ability under conditions in which this residue was charged, that is, where the hydrophobicity of the 8^th^ residue was much lower than the hypothesized required range (Figure 3). Furthermore, by analyzing alignments of native TatA sequences, we found that species from different kingdoms have different preferred amino acids at the 8^th^ position correlating with conserved 4-residue motifs at the N-terminus of TMH (Figure 5). Such 4-residue motifs were further tested in *E. coli* TatA and proved to have significant effect on the transport ability (Figure 6). Interestingly, these motifs were correlated with the predominance of usage of the ΔpH or Δψ components of the pmf (Figure 7). In the congregate, these results provide new perspectives through which to understand the functional significance of the 8^th^ residue in TatAs.

Our finding that the hydrophobicity, but not polarity, of the 8^th^ amino acid, is critical for TatA supports the idea that this residue’s ability to respond to membrane potential or to electrically interact with other polar amino acids in other Tat subunits is not necessary for the fundamental transport mechanism. Given that *E. coli* TatA lacks a charged polar amino acid whilst chloroplast TatAs have a charged residue at the corresponding position, the possibility that bacteria and thylakoids might use different mechanisms in terms of TatA function has been discussed (New et al., 2018). It has been debated whether an active shift of protonation states of the glutamic acid in the chloroplasts TatA is required for a functional transporter (Berks, 2015; New et al., 2018). However, this cannot be the case since many TatAs do not have this protonatable residue, and we find that even Ala in this position confers some activity. Instead, the requirement appears to be for a residue with a moderate hydrophobicity and helix stability. In that context, the pKa of the side chain of glutamic acid matches the thylakoid’s lumen pH in the light (Kramer et al., 1999), and this changes the overall hydrophobicity of the amino acid (Kovacs et al., 2006) to the moderate range. Such an explanation is further supported by two observations. First, it has been shown that thylakoid Tat transport operates with a delay between the light illumination and transport event (a sigmoidal transport curve), even at saturating Tha4 concentrations (Celedon and Cline, 2012). Such a delay cannot be completely attributed to the translocon assembly step, and may also be caused by the time required to develop a lumen pH low enough to modulate the Glu hydrophobicity. Second, our sequence alignments revealed that nine archaea species have TatAs with an aspartic acid (Supplemental Figure 3E) at the corresponding position whose side chain’s pKa is even lower than that of glutamic acid. Interestingly, we found by analyzing the living environments for those species that all of them live in extreme acidic environments (Supplemental Table 1). A random sample of archaea species with E8 did not reveal any acidic extremophiles. While these observations do not explain why the TatAs have acidic amino acids that require neutralizing in the first place, they do provide examples of how Tat system might be altered to accommodate to different physiological conditions.

Our experiments suggest that the creation of a relatively unstable N-terminus structure by placing a polar 8^th^ residue within the hydrophobic TatA TMH is likely significant. Such a residue would be expected to be located at or near the membrane/water interface, thereby biasing the position the N-terminus of the TMH close to the membrane surface(Pettersson et al., 2018). Meanwhile, a recent study reported that the *E. coli* TatA TMH is tuned to be 15-residues long to keep the hydrophobic mismatch while also maintaining membrane integrity (Hao et al., 2021; Mehner-Breitfeld et al., 2021) Also, it has been shown that TatAs have a highly conserved 12-residue hydrophobic core (9^th^-20^th^ in *E. coli* TatA) in the TMH (Hao et al., 2021; Mehner-Breitfeld et al., 2021). Together these observations indicate that the TatA TMH could be as short as 12 amino acids while the 8^th^ residue is transiently exposed to the aqueous environment. This might allow membrane bilayer rupture (i.e., toroidal pore formation), with concomitant transport of a substrate held at the site. Our data herein demonstrated that if the 8^th^ residue was too hydrophobic (Figure 1E), Tat transport was completely blocked, even though the hydrophobic mismatch between TatA TMH and membrane bilayer was still maintained. This suggests that a 15-residue long TMH is required but insufficient for active Tat transport, and a transition from that length to an even shorter TMH might be functionally important.

We showed that not only the 8^th^ residue is critical, but also found a changing four-residue motif at the N-terminus of the TatA TMH affects the overall TatA function (Figure 5 and 6). This motif was perhaps not identified in previous studies for two reasons. First, the functional N-terminus motif was not obviously conserved across different species. For example, in chloroplasts and cyanobacteria, a “GxPE” motif was nearly invariant at the N-terminus of the TatA TMH. However, this motif is not present in proteobacterial species. Second, the identified N-terminal motifs showed strong correlation with the 8^th^ residue in the TatA TMH. For instance, in proteobacteria, “SIW” was conserved in the 5^th^ to 7^th^ position when the 8^th^ residue was glutamine or histidine, whereas this motif was not conserved when lysine or glycine was at the 8^th^ position (Supplemental Figure 3). Even though we still do not understand the explicit roles of the conserved motifs, the amino acid choices therein likely provide clues about the function of N-terminus of the TatA TMH. For instance, a tryptophan was observed immediately before H8 or Q8 in proteobacteria and in two groups of gram-positive bacteria. One question would be what are the special features of tryptophan that cause it to be conserved in this context. Tryptophan is normally considered to be a non-polar amino acid in biochemistry textbooks. However, it possesses a highly polarizable indole group in the side chain. This results in tryptophan being highly abundant at the water-lipid interface and acting as an anchor to stabilize membrane protein segments (de Jesus and Allen, 2013; Sun et al., 2008). Moreover, the indole group could respond to an electric field and lower the conformational energy of the TMH were it to algin with the electrical force lines (Zhang and Li, 2019). Thus, the tryptophan might help TatA embed in the membrane and serve as an anchor near the interface. Additionally, we noticed that a proline residue is highly conserved at position 7 when a glutamic acid is in position 8 in TatAs. Proline is generally considered a helix breaker in soluble proteins (Barlow and Thornton, 1988). However, it has been suggested that proline would not disrupt an alpha-helix in a membrane environment (Li et al., 1996; Senes et al., 2004).

Moreover, it has been shown that the hinge-bending motions of proline residue are likely to play a significant role in catalysis and signal transduction (S.P. Sansom and Weinstein, 2000; Tieleman et al., 2001). Such a hinge forming function at the N-terminus of the TatA TMH is consistent with our hypothesis that a transition to an even shorter TMH might be beneficial for the Tat mechanism. In addition, according to the TatA NMR structure (Hu et al., 2010; Pettersson et al., 2018), this 4-residue motif presents in the first coil of the TMH. Ser6 and Gln8 orientate oppositely than Ile6 and Trp7, suggesting a weak local amphipathic topology. Moreover, other 4-residues motifs in chloroplasts and gram-positive bacteria are also arrayed this way, even though they have different combinations of four residues at those positions. It is possible that this local amphipathic character assists in the reported motion of the TMH into and out of the membrane during transport (Aldridge et al., 2012).

Incorporating our new findings related to the function of TatA subunits we propose a speculative new model for the mechanism of Tat pathway (Figure 8). In this model, presented here to stimulate discussion: (i) Before substrate binding, TatA, TatB and TatC form an oligomeric structure which serves as the receptor complex for signal peptide of Tat substrates (Alcock et al., 2016). At this stage, relatively few TatA proteins are associated with the TatBC complex. (ii) – (iii) After substrate binding and in the presence of a pmf, the receptor complex undergoes a conformation change that attracts more TatA proteins (Aldridge et al., 2014). Unlike previous models (Rodriguez et al., 2013), we take the view of (Mehner-Breitfeld et al., 2021) and propose these additional TatA proteins will be initially recruited into a non-circular structure, perhaps arrayed as the double line illustrated. (iv) As more TatA proteins are recruited to the translocon, the hydrophobic mismatch, unstable N-terminus structure, and the membrane-thinning action of the pmf (Matthew P Johnson et al., 2011; Matthew P. Johnson et al., 2011; Kirchhoff et al., 2011; Murakami and Packer, 1970) cause the double line to separate and join with subunits in adjacent lines to form a ring with a tight-fitting toroidal pore around the substrate (v). (vi) The concomitant movement of protons during substrate translocation (Alder and Theg, 2003) results in a local dissipation of the pmf, and this in turn returns the membrane to a thickness that favors the bilayer, causing dispersal of TatA from the translocation site and return of the translocon to its resting condition. While certain aspects of this model are admittedly speculative, we believe it captures many essential properties of Tat transport discerned from experiments. Future studies probing the biophysical properties of the membranes at the site of protein translocation are likely to shed further light on the mechanism of this enigmatic protein transport system.

**Figure 8.**
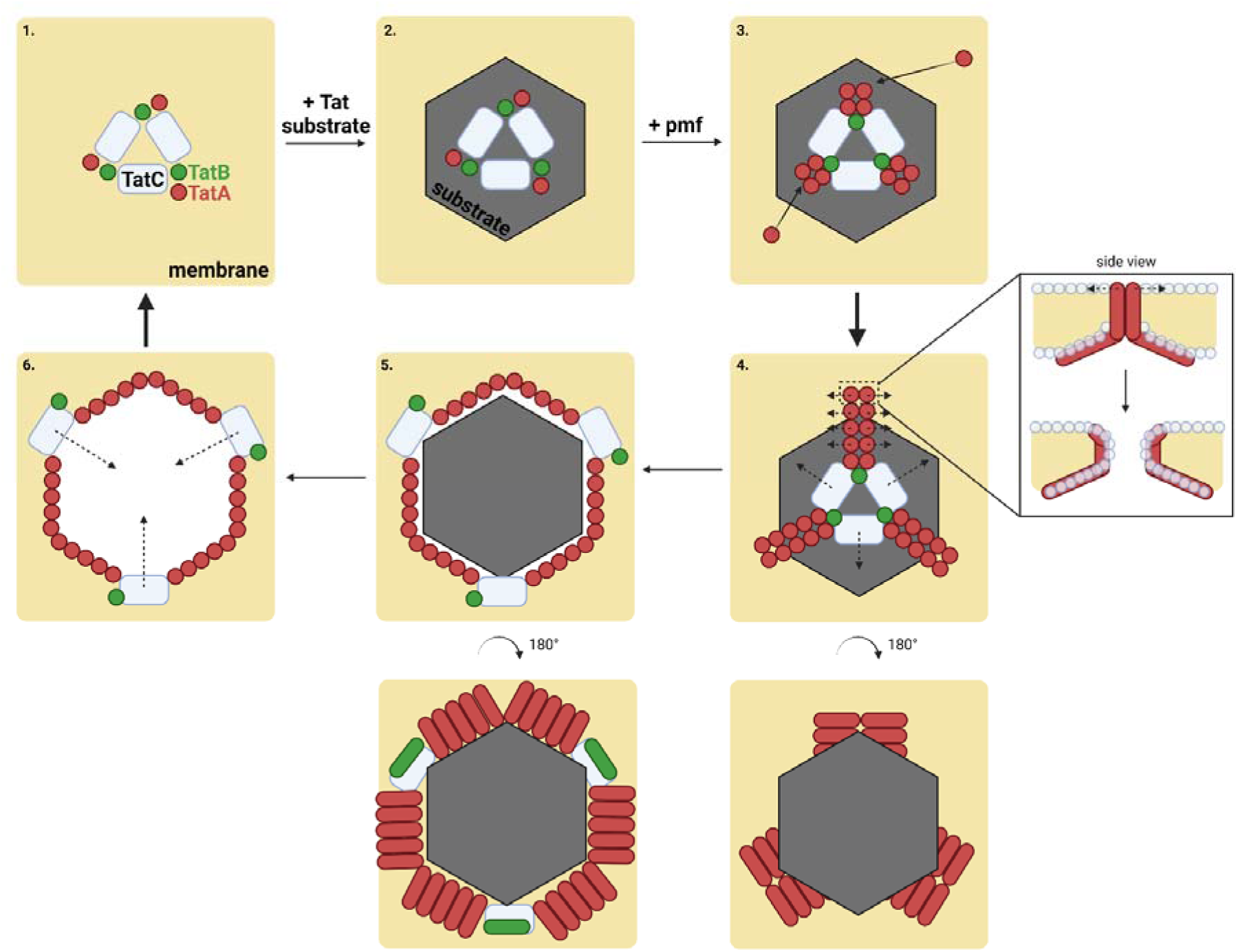
Schematic diagram for the proposed Tat transport mechanism. In this diagram, a view of the proposed Tat transport process is shown from above the membrane. Side view of the TatA interfaces is shown on the right of the step 4. Blue/grey circles represent lipid head group.

## Methods

### 1. Strain and Plasmid Constructions

*E. coli* strain DADE-A (MC4100, Δ*tatABCE*, arabinose resistance) was used in this study (Wexler et al., 2000). For TatA variants in pTat101 (Kneuper et al., 2012), which is a low copy plasmid expressing the TatABC constitutively, the corresponding single and multiple amino acid substitutions were introduced by QuikChange site-directed mutagenesis (NEB, Phusion High-Fidelity PCR Kit) followed by KLD (Kinase, Ligase, and DpnI) digestion (NEB). Detailed primers used for TatA variant construction can be found in Supplemental Table 2. For TatA variants in pBAD22, which is a vector with an arabinose-inducible *araBAD* operon, indicated TatA alleles, together with the wild-type TatBC, were amplified from the respective constructs in pTat101. For TatA single Q8 substitution, the forward primer TatA_gopBAD22_F (5’-ACC ACA GAG GAA CAT GTA TGG GTG GTA TCA GTA TTT G-3’) was used. For TatA multiple substitutions, the forward primer TatAnmotif_gopBAD22_F (5’- CTA CCA CAG AGG AAC ATG TAT GGG TGG TAT C-3’) was used. Reverse primer TatABC_gopBAD22_R (5’-TCT AGA GGC GGT TGA ATT TAT TCT TCA GTT TTT TCG C-3’) was used in both cases. Empty vectors were constructed using the primers pBAD22delABC_F (5’-ATT CAA CCG CCT CTA GAG-3’) and pBAD22delABC_R (5’-ACA TGT TCC TCT GTG GTA G-3’). Fragments and vector were assembled using the Gibson Assembly approach (NEBuilder HiFi DNA Assembly Master Mix). For TatA variants in pBAD33, which is a vector with an arabinose-inducible *araBAD* operon, a 6X His tag was first added at the C-terminus of TatA by using the primers pBAD33His_F (5’- CATCATCATCACCACCACTAATGGC-3’) and pBAD33His_R (5’-TGATGATGACCCACCTGCTCTTTATCGTG-3’). Subsequently, TatA mutations were introduced by QuikChange site-directed mutagenesis as described above. pQE80l (SufI-FLAG) for the *in vivo* transport assay was constructed as describe previously (Huang and Palmer, 2017). pNR14 and pNR42, two plasmids used in pulse chase experiments, were as described previously (Sargent et al., 1999; Stanley et al., 2000). All plasmids were confirmed by Sanger sequencing. More detailed information on plasmid constructions can be found in Supplemental Table 3.

### 2. Sequence Alignments and Statistical Analyses

A total of 15054 TatA sequences were downloaded from the identical protein groups in GenBank (NCBI). Sequences were then divided into five sub-groups based on taxonomy (7534 from proteobacteria, 6978 from firmicutes and actinobacteria, 149 from cyanobacteria, 242 from chloroplast, and 151 from archaea), and were subsequently subjected to multiple sequence alignments within each group using MAFFT (Rozewicki et al., 2019). Sequence logos corresponding to each group were generated using RStudio (Ver. 1.4.1103) with ggseqlogo package (Wagih, 2017). For the interest of this project, sequences were further grouped by the identity of the residue aligned to the eighth position in *E. coli* TatA, and the corresponding sequence logos were generated. A stacked bar chart summarizing the distribution of the identity of residues in this position in each taxonomical group was plotted using GraphPad Prism version 8.2.1 for Windows (GraphPad Software, San Diego, California USA).

### 3. Liquid SDS Growth Assay

Overnight bacteria cultures were sub-cultured with an optical density (OD_600_) of 0.002 were grown in the Luria-Bertani (LB) medium containing 0%, 5%, or 10% SDS, respectively. After cells were incubated at 37°C with shaking for 5 hours, the final OD_600_ were measured. For each mutant, survival ratio was calculated by its final OD_600_ at the indicated SDS concentration to the final OD_600_ in the LB medium without SDS.

### 4. *in vivo* Transport Assay and Cell Fractionation

Corresponding mutant plasmids derived from pTat101 were co-transformed with pQE80l (SufI-FLAG), which under the control of the T5 promoter, into the DADE-A strain. Overnight cultures were diluted in a fresh LB medium at a ratio of 1:100 and grown at 37 °C until the OD_600_ reached 0.5. A final concentration of 1 mM IPTG (isopropyl β-D-1-thiogalactopyranoside) was added to induce the expression of SufI-FLAG. After cultured at 37°C for another 2.5 hours, 3 mL of the cells with an OD_600_ equivalent to 1 were fractionated using the EDTA/lysozyme/cold osmotic shock method described before (Petiti et al., 2017b). Periplasmic fractions were collected by adding equal volume of 2X SDS sample buffer, which were then subjected to SDS-PAGE. Membrane fractions were washed in 10 mM Na_2_CO_3_ for 1 hour at 4°C with gentle shaking. Subsequently, samples were ultracentrifuged at 120,000 x g for 45 min at 4°C. Pellets were resuspended with 2X SDS sample buffer and collected as the membrane fractions.

### 5. SDS-PAGE and Western Blot

For periplasmic fractions, samples were immunoblotted with α-FLAG (GenScript) followed by HRP-conjugated α-mouse antibodies (Santa Cruz Biotechnology). For membrane fractions, after transfer the proteins to the PVDF membranes, PVDF membranes were cut and immunoblotted with α-TatA and α-TatB antibodies respectively followed by HRP-conjugated α-rabbit antibody (GenScript). Proteins were visualized using ProSignal Pico ECL Western Blotting detection kit (Genesee Scientific).

### 6. Pulse-Chase Experiment and Autoradiography

The pulse-chase experiment protocol was modified from (Stanley et al., 2000). Tat variants carrying pNR14 and pNR42 were grown overnight in LB media at 30°C, which were then sub-cultured in fresh LB media the next day for 1.5 hours at 30°C. Subsequently, cells were harvested and normalized to the equivalent of 0.5 mL cells with OD_600_ = 0.2. Cells were washed with 0.5 mL 1X M9 medium (diluted from 10X M9 medium supplemented with 0.1 mM CaCl_2_, 0.002% thiamine, 2 mM MgSO_4_, and 0.01% of the 18 amino acids excluding methionine and cysteine, pH = 7.0). Cells were resuspended in 2.5 mL M9 medium and cultured for another hour at 30°C. Then, cells were transferred to 42°C for 15 min to induce T7 polymerase from pNR42. 400 μg/mL of rifampicin was added to inhibit the *E. coli* endogenous RNA polymerase, and cells were grown at 42°C for another 10 min. Subsequently, cells were grown at 30°C for another 20 mins before transferred to the indicated temperature until the completion of the experiment. If needed, pH was adjusted by 1 M HCl or 1 M KOH immediately after this step. Bacteria cultures were then incubated at indicated temperature for 5 min. 0.025 mCi of [^35^S] methionine (PerkinElmer Inc. NEG772002MC) was added for the pulse step, and 750 μg/mL of unlabeled cold methionine was added after 5 min. 300 μL of the culture was taken at the indicated time point, and transport was quenched by freezing in liquid nitrogen. Samples were subsequently thawed on ice, centrifuge, and resuspended with 2X SDS sample buffer followed by SDS-PAGE and autoradiography. Results were quantified using ImageJ software.

### 7. Statistical Analysis of the Pulse-Chase Experimental Results

Mature-to-total (sum of precursor and mature) data from the pulse-chase experiment was plotted against each time point, which were then fitted with the exponential plateau model deriving from the first-order reaction model, *y* = (*y*_max_ – *y*_0_) × *e*^−*kt*^, using GraphPad Prism. *y*_max_ was defined as the maximum of the mature-to-total value, and *y*_0_ represents the initial mature-to-total value at the end of the pulse. t represents time in minutes. the initial velocity (*V*_0_), representing the transport rate when *y* = 0, was obtained by *V*_0_ = *y*_max_ × *k*, with the units of the reciprocal minutes.

### 8. IMVs (Inverted-Membrane Vesicles) preparation

Tat proteins expression and IMVs preparation were performed as described (Bageshwar and Musser, 2007b), with the following modifications. Cells were resuspended with the same lysis buffer as described, which were then passed through a French press at ∼ 6000 psi once. Subsequently, cells were centrifuged at 30,000 g for 20 mins at 4°C. Supernatants were then centrifuged at 140,000 g for 1 hr. Pellets were resuspended with IMV storage buffer (10 mM Tris-HCl, pH = 7.5, 1 mM MgSO_4_, 1 mM KCl, 40% glycerol). IMVs were stored in -80°C freezer.

### 9. Proton Leakage Measurement

IMVs were made from the DADE-A cells expressing the 6X His-tagged wild-type or TatA variants in pBAD33 alone or together with the pTat101(Δ*tatA*). Acridine orange fluorescence-quenching assays were carried out using the Fluorolog-3 spectrofluorometer (HORIBA Scientific, model No. FL3-22). IMVs were added to the final concentration equivalent to A_280_ = 0.375 to the reaction mix containing 1X TE buffer (25 mM MOPS, 25 mM MES, 5 mM MgCl_2_, 50 mM KCl, 200 mM sucrose, and 57 μg/mL BSA, pH = 7.0), ATP regeneration system (2.9 mM phosphocreatine, 0.29 mg/mL creatine kinase), and 2 μM acridine orange. The reaction mix was incubated for 5 mins with gentle stirring at 37°C before measurement. The fluorescence of acridine orange was recorded at λ_ex_ = 494 nm (slit = 1 nm) and λ_em_ = 540 nm (slit = 5 nm) per 0.1 second. 4 mM of ATP was added at 60 seconds, and 10 μM of CCCP was added at 200 seconds to dissipate the proton gradient.

## Acknowledgement

We gratefully acknowledge support from the Division of Chemical Sciences, Geosciences, and Biosciences, Office of Basic Energy Sciences of the U.S. Department of Energy through Grant DE-SC0020304 to SMT.

