## Supplemental Information for "Functional analysis of the polar amino acid in the TatA transmembrane helix"

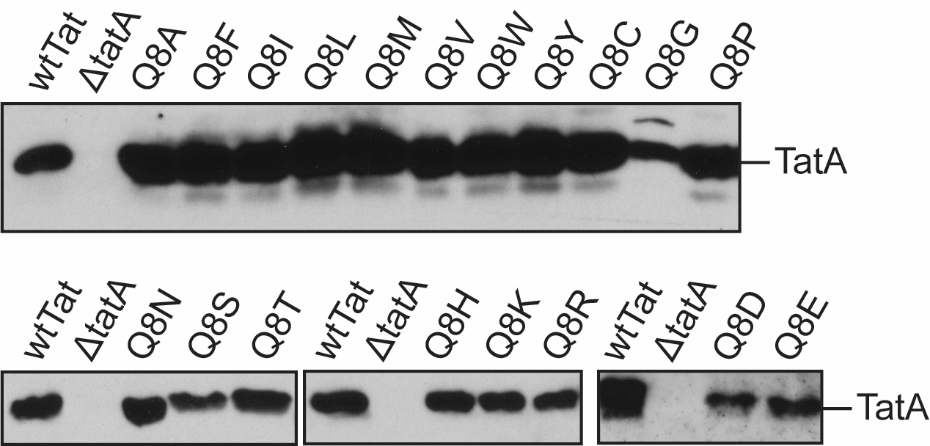

**Supplemental Figure 1. Membrane stability of the TatA Q8 substituted mutants.** The membrane stability of the TatA Q8 mutants together with the wtTatA and *ΔtatA* were assessed by the immunoblotting of the carbonate-treated membrane fraction from whole-cell extracts. Anti-TatA antibody was used to detect the presence of TatA in the membrane fractions.

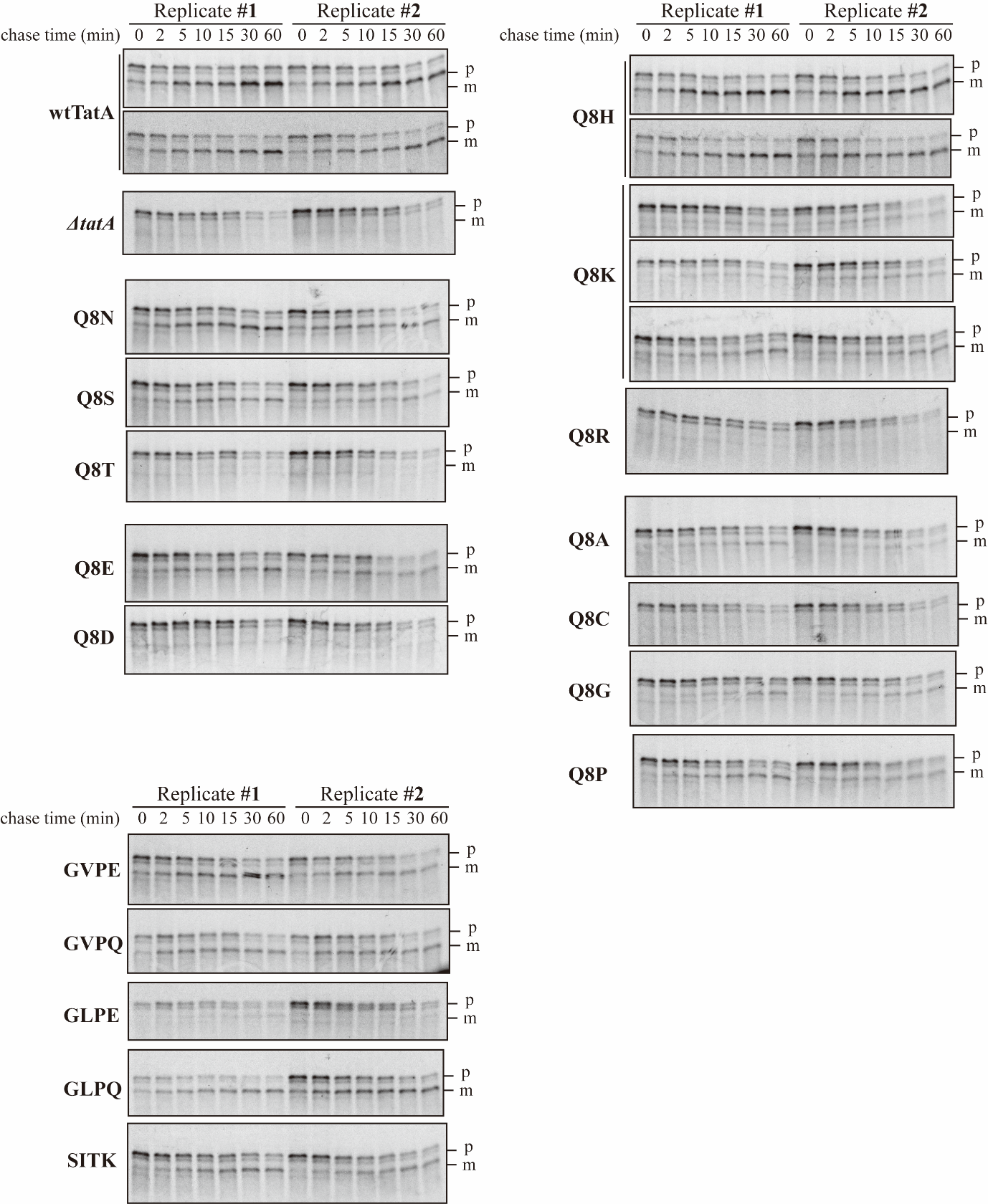

**Supplemental figure 2. The autoradiography results of pulse-chase experiments.** At least two replicates were performed for each mutant. Identities of the mutant is shown on the left. Replicate number and the corresponding time of chase step are shown on the top. Samples were subjected to SDS-PAGE using 8-16% polyacrylamide gels. p, precursor; m, mature.

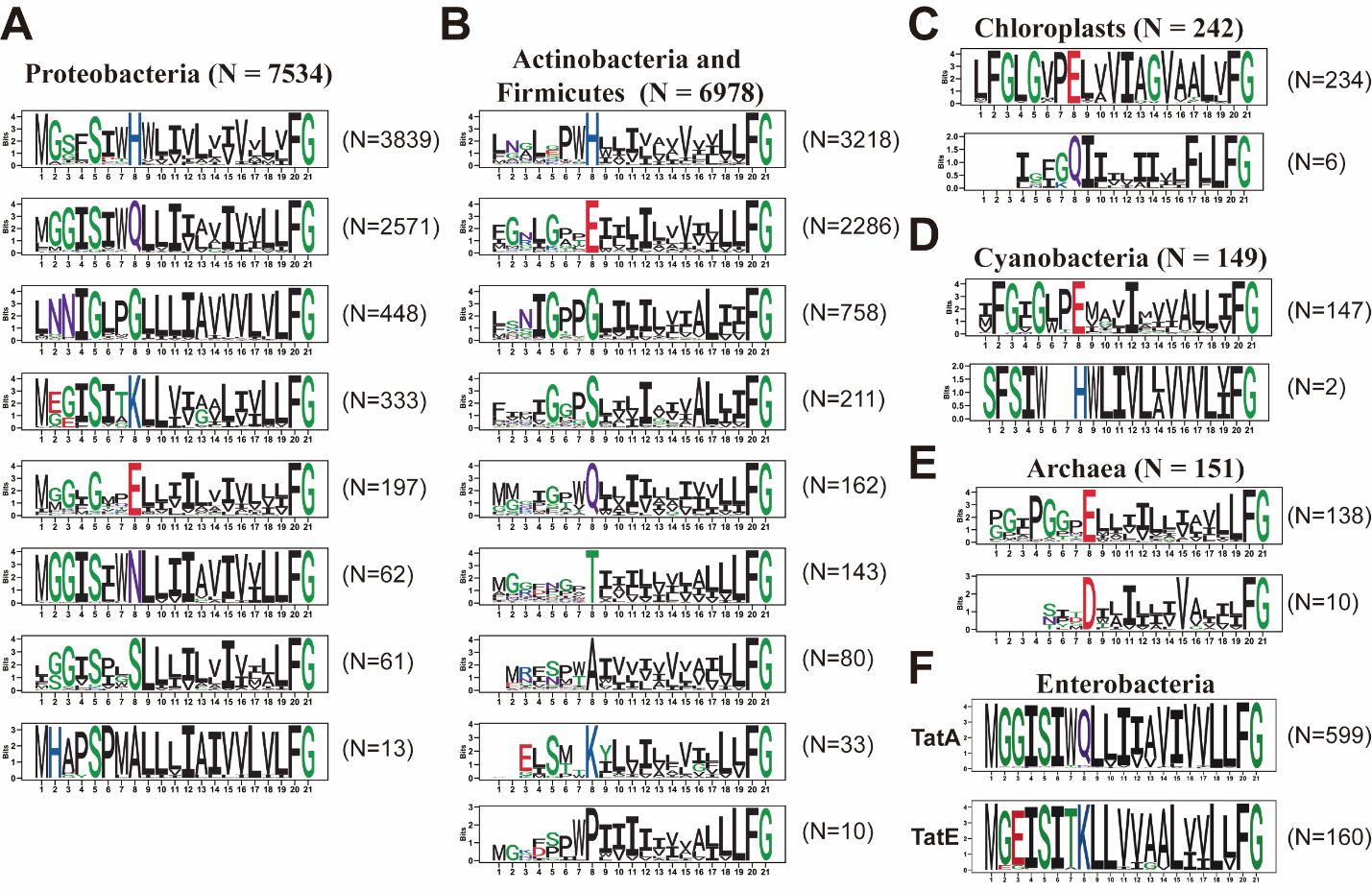
**Supplemental figure 3. Detailed sequence logos across different species.** Alignment data were subset based on the identity of the amino acid at the position equivalent to the eighth one in the *E. coli* TatA. N, number of sequences used to generate the sequence logo. (A) Sequence logos for proteobacteria. (B) Sequence logos for actinobacteria and firmicutes. (C) Sequence logos for chloroplasts. (D) Sequence logos for cyanobacteria. (E) Sequence logos for archaea. (F) A subset of data, enterobacteria, which possess TatE, were separated from proteobacteria to generate the sequence logos.

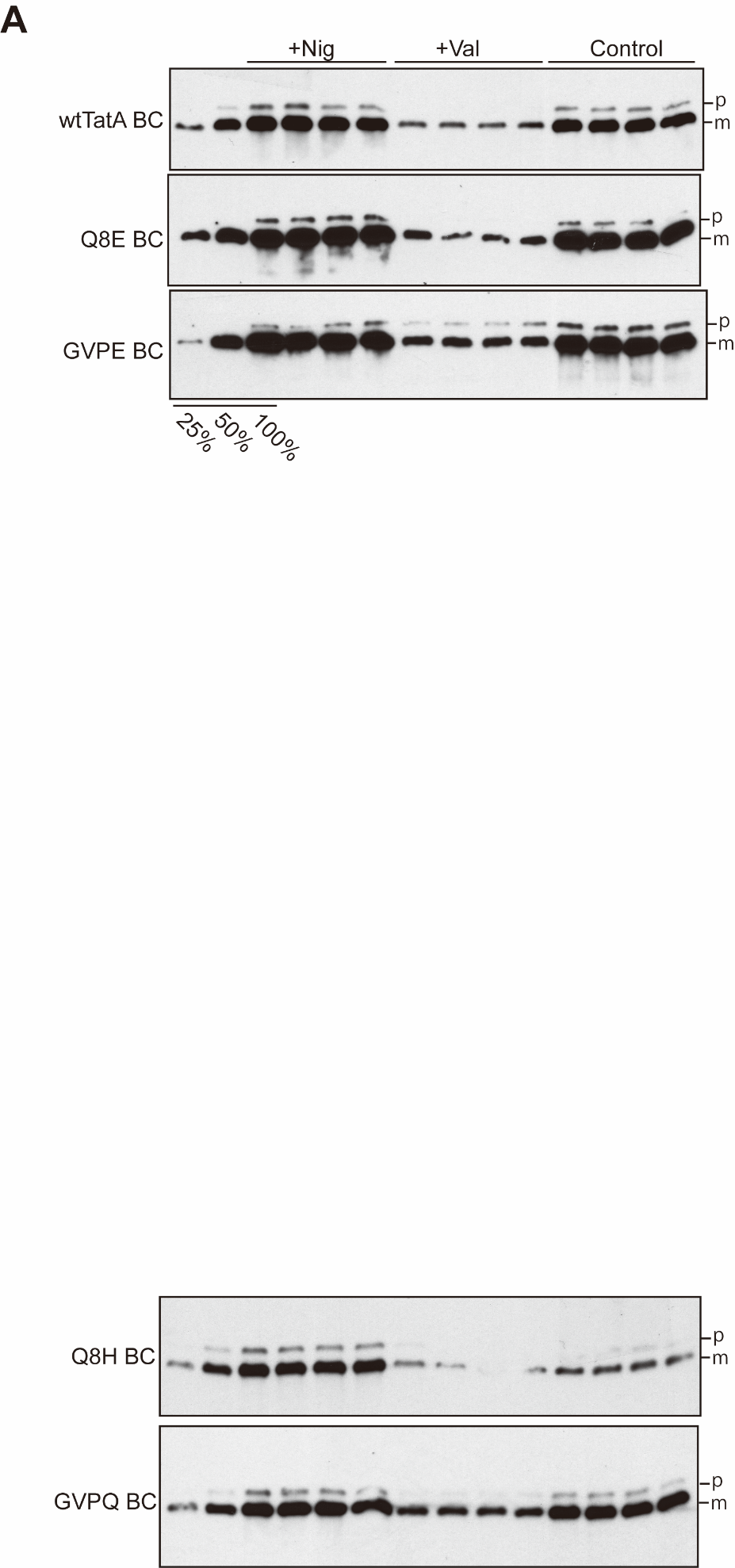

**Supplemental figure 4. Western-blot results of the IMV transport assay.** IMVs transporting biotinylated SufI were subjected to proteinase K digestion to remove the non-transported SufI. Digested samples were subjected to SDS-PAGE using 8-16% polyacrylamide gels followed by Western-blot using avidin to detect the biotinylated SufI. Identities of the IMVs are shown to the left. Experimental treatments are shown on the top. p, precursor; m, mature. Four replicates were included in this experiment.

**Supplemental Table 1**

| Species name | Residue at 8th position | Living condition | Data source |
| --- | --- | --- | --- |
| *Acidithrix ferrooxidans* | Asp (D) | pH 1.5 | (Jones and Johnson, 2015) |
| *Corynebacterium sp. DSM 45110* | Asp (D) | n.d. | N/A |
| *Candidatus Aramenus sulfurataquae* | Asp (D) | Muddy water from acid hot spring | The data have been deposited with links to BioProject accession number PRJNA273893 in the NCBI BioProject database (https://www.ncbi.nlm.nih.gov/bioproject/). |
| *Ferroplasma acidarmanus* | Asp (D) | pH 0 - 2.5 | (Baumler et al., 2005) |
| *Picrophilus torridus* | Asp (D) | pH 0 | (Fütterer et al., 2004) |
| *Metallosphaera cuprina* | Asp (D) | pH 3.5, 55 - 75°C | (Liu et al., n.d.) |
| *Ferroplasma acidiphilum* | Asp (D) | pH 1.7 | (Golyshina et al., n.d.) |
| *Picrophilus oshimae* | Asp (D) | pH < 3.5, optimal pH 0.7 | (Schleper et al., 1995) |
| *Sulfurisphaera ohwakuensis* | Asp (D) | pH 1.0 - 5.0, optimal pH 2.0 | (Kurosawa et al., n.d.) |

n.d. = no data; N/A = not applicable.

**Supplemental Table 2. Primer sequences in this study.**

| **Primer Name** | **Primer Sequence** |
| --- | --- |
| TatA_Q8A_F | 5’-CAGTATTTGGgcgTTATTGATTATTGCCGTC-3’ |
| TatA_Q8C_F | 5’-CAGTATTTGGtgcTTATTGATTATTGCCGTCATCGTTG-3’ |
| TatA_Q8D_F | 5’-CAGTATTTGGgatTTATTGATTATTGCCG -3’ |
| TatA_Q8E_F | 5’-CAGTATTTGGgagTTATTGATTATTGC-3’ |
| TatA_Q8F_F | 5’-CAGTATTTGGtttTTATTGATTATTGCCGTCATCGTTG-3’ |
| TatA_Q8G_F | 5’-CAGTATTTGGggtTTATTGATTATTGCCGTCATCG-3’ |
| TatA_Q8H_F | 5’-CAGTATTTGGcacTTATTGATTATTGCCG-3’ |
| TatA_Q8I_F | 5’-CAGTATTTGGattTTATTGATTATTGCCGTCATCG-3’ |
| TatA_Q8K_F | 5’-CAGTATTTGGaagTTATTGATTATTGC-3’ |
| TatA_Q8L_F | 5’-CAGTATTTGGctgTTATTGATTATTGC-3’ |
| TatA_Q8M_F | 5’-CAGTATTTGGatgTTATTGATTATTGCCG-3’ |
| TatA_Q8N_F | 5’-CAGTATTTGGaatTTATTGATTATTGCCGTC-3’ |
| TatA_Q8P_F | 5’-CAGTATTTGGccgTTATTGATTATTGC-3’ |
| TatA_Q8R_F | 5’-CAGTATTTGGagaTTATTGATTATTGCCGTCATCG-3’ |
| TatA_Q8S_F | 5’-CAGTATTTGGagcTTATTGATTATTGCCGTC-3’ |
| TatA_Q8T_F | 5’-CAGTATTTGGaccTTATTGATTATTGCCGTCATCG-3’ |
| TatA_Q8V_F | 5’-CAGTATTTGGgtgTTATTGATTATTGCCGTC-3’ |
| TatA_Q8W_F | 5’-CAGTATTTGGtggTTATTGATTATTGCCGTC-3’ |
| TatA_Q8Y_F | 5’-CAGTATTTGGtatTTATTGATTATTGCCGTC-3’ |
| TatA_Q8_R | 5’-ATACCACCCATGGATCCTC-3’ |
| TatA_GLPE_F | 5’-cctggaTTATTGATTATTGCCGTCATC-3’ |
| TatA_GLPE_R | 5’-aagaccGATACCACCCATGGATCC-3’ |
| TatA_GLPQ_F | 5’-accgCAGTTATTGATTATTGCCGTC-3’ |
| TatA_GLPQ_R | 5’-aaaccGATACCACCCATGGATCC-3’ |
| TatA_GVPE_F | 5’-ccggagTTATTGATTATTGCCGTCATC-3’ |
| TatA_GVPE_R | 5’-aacaccGATACCACCCATGGATCC-3’ |
| TatA_GVPQ_F | 5’-ccgcagTTATTGATTATTGCCGTCATC-3’ |
| TatA_GVPQ_R | 5’-aacaccGATACCACCCATGGATCC-3’ |
| TatA_SITK_F | 5’-accaaaTTATTGATTATTGCCGTCATC-3’ |
| TatA_SITK_R | 5’-aatactGATACCACCCATGGATCC-3’ |
| pBAD33(TatA_Q8E)_F | 5’-CAGTATTTGGgagTTATTGATTATTG-3’ |
| pBAD33(TatA_Q8K)_F | 5’-CAGTATTTGGaaaTTATTGATTATTGC-3’ |
| pBAD33(TatA_Q8)_R | 5’-ATACCACCCATGACCTTTC-3’ |
| pBAD33(TatA_7P8E)_F | 5’-ccggagTTATTGATTATTGCCGTCATC-3’ |
| pBAD33(TatA_7P8Q)_F | 5’-ccgcagTTATTGATTATTGCCGTCATC-3’ |
| pBAD33(TatA_5G6V)_R | 5’-cacgccGATACCACCCATGACCTTTC-3’ |
| pBAD33(TatA_5G6L)_F | 5’-caggccGATACCACCCATGACCTTTC -3’ |

Note: mutagenesis sites are shown in lowercase.

**Supplemental Table 3. Plasmids information in this study.**

| Name | Description | Source |
| --- | --- | --- |
| **Vector** | | |
| pBAD22 | Expression vector with arabinose-inducible *araBAD* operon, carbenicillin resistance | ATCC |
| pBAD33 | Expression vector with arabinose-inducible *araBAD* operon, chloramphenicol resistance | (Guzman et al., 1995) |
| pQE80l | Expression vector with IPTG-inducible T5 promoter | Qiagen |
| **Substrate** | | |
| pNR14 | *E. coli* SufI in pT7.5 vector with T7 φ10 promoter | (Stanley et al., 2000) |
| pNR42 | Phage T7 polymerase and the temperature-sensitive λ repressor  in pSU18 | (Sargent et al., 1999) |
| pQE80l (SufI-FLAG) | *E. coli* SufI with C-terminal FLAG tag in pQE80l vector | This work |
| **Mutant** | | |
| pTat101 | pTH19Kr derivative, a low copy vector, expression of TatABC | (Kneuper et al., 2012) |
| ΔTatA | pTH19Kr derivative, a low copy vector, expression of TatBC | This work |
| pTat101 (Q8A) | pTat101 with Ala substitution for Gln8 in TatA | This work |
| pTat101 (Q8C) | pTat101 with Cys substitution for Gln8 in TatA | This work |
| pTat101 (Q8D) | pTat101 with Asp substitution for Gln8 in TatA | This work |
| pTat101 (Q8E) | pTat101 with Glu substitution for Gln8 in TatA | This work |
| pTat101 (Q8F) | pTat101 with Phe substitution for Gln8 in TatA | This work |
| pTat101 (Q8G) | pTat101 with Gly substitution for Gln8 in TatA | This work |
| pTat101 (Q8H) | pTat101 with His substitution for Gln8 in TatA | This work |
| pTat101 (Q8I) | pTat101 with Ile substitution for Gln8 in TatA | This work |
| pTat101 (Q8K) | pTat101 with Lys substitution for Gln8 in TatA | This work |
| pTat101 (Q8L) | pTat101 with Leu substitution for Gln8 in TatA | This work |
| pTat101 (Q8M) | pTat101 with Met substitution for Gln8 in TatA | This work |
| pTat101 (Q8N) | pTat101 with Asn substitution for Gln8 in TatA | This work |
| pTat101 (Q8P) | pTat101 with Pro substitution for Gln8 in TatA | This work |
| pTat101 (Q8R) | pTat101 with Arg substitution for Gln8 in TatA | This work |
| pTat101 (Q8S) | pTat101 with Ser substitution for Gln8 in TatA | This work |
| pTat101 (Q8T) | pTat101 with Thr substitution for Gln8 in TatA | This work |
| pTat101 (Q8V) | pTat101 with Val substitution for Gln8 in TatA | This work |
| pTat101 (Q8W) | pTat101 with Trp substitution for Gln8 in TatA | This work |
| pTat101 (Q8Y) | pTat101 with Tyr substitution for Gln8 in TatA | This work |
| pTat101 (Ad2) | pTat101 with Leu18 and Leu19 deletion in TatA |  |
| pTat101 (GLPE) | pTat101 with Gly substitution for Ser5, Leu substitution for Ile6, Pro substitution for Trp7, and Glu substitution for Gln8 in TatA | This work |
| pTat101 (GLPQ) | pTat101 with Gly substitution for Ser5, Leu substitution for Ile6, Pro substitution for Trp7 in TatA | This work |
| pTat101 (GVPE) | pTat101 with Gly substitution for Ser5, Val substitution for Ile6, Pro substitution for Trp7, and Glu substitution for Gln8 in TatA | This work |
| pTat101 (GVPQ) | pTat101 with Gly substitution for Ser5, Val substitution for Ile6, Pro substitution for Trp7 in TatA | This work |
| pTat101 (SITK) | pTat101 with Thr substitution for Trp7, and Lys substitution for Gln8 in TatA | This work |
| pTat101 (GVPE(ACa1)) | pTat101 (GVPE) with one Leu addition after Leu19 in TatA | This work |
| pTat101 (GVPE(ACa2)) | pTat101 (GVPE) with two Leu additions after Leu19 in TatA | This work |
| pTat101 (GVPQ(ACa1)) | pTat101 (GVPQ) with one Leu addition after Leu19 in TatA | This work |
| pTat101 (GVPQ(ACa2)) | pTat101 (GVPQ) with two Leu additions after Leu19 in TatA | This work |
| pBAD22(TatABC) | TatABC in pBAD22 | This work |
| pBAD22 (Q8E) | TatA with Glu substitution for Gln8 in pBAD22 | This work |
| pBAD22 (Q8K) | TatA with Lys substitution for Gln8 in pBAD22 | This work |
| pBAD22 (GLPE) | TatA with Gly substitution for Ser5, Leu substitution for Ile6, Pro substitution for Trp7, and Glu substitution for Gln8 in pBAD22 | This work |
| pBAD22 (GLPQ) | TatA with Gly substitution for Ser5, Leu substitution for Ile6, Pro substitution for Trp7 in pBAD22 | This work |
| pBAD22 (GVPE) | TatA with Gly substitution for Ser5, Val substitution for Ile6, Pro substitution for Trp7, and Glu substitution for Gln8 in pBAD22 | This work |
| pBAD22 (GVPQ) | TatA with Gly substitution for Ser5, Val substitution for Ile6, Pro substitution for Trp7 in pBAD22 | This work |
| pBAD33(TatAhis) | TatA with a C-terminal 6X His in pBAD33 | This work |
| pBAD33 (Q8Ehis) | TatA with Glu substitution for Gln8 and a C-terminal 6X His in pBAD33 | This work |
| pBAD33 (Q8Khis) | TatA with Lys substitution for Gln8 and a C-terminal 6X His in pBAD33 | This work |
| pBAD33 (GLPEhis) | TatA with Gly substitution for Ser5, Leu substitution for Ile6, Pro substitution for Trp7, Glu substitution for Gln8, and a C-terminal 6X His in pBAD33 | This work |
| pBAD33 (GLPQhis) | TatA with Gly substitution for Ser5, Leu substitution for Ile6, Pro substitution for Trp7, and a C-terminal 6X His in pBAD33 | This work |
| pBAD33 (GVPEhis) | TatA with Gly substitution for Ser5, Val substitution for Ile6, Pro substitution for Trp7, Glu substitution for Gln8, and a C-terminal 6X His in pBAD33 | This work |
| pBAD33 (GVPQhis) | TatA with Gly substitution for Ser5, Val substitution for Ile6, Pro substitution for Trp7, and a C-terminal 6X His in pBAD33 | This work |
